# A Large-scale Study Reveals 24 hour Operational Rhythms in Hospital Treatment

**DOI:** 10.1101/617944

**Authors:** Marc D. Ruben, Lauren J. Francey, Yuping Guo, Gang Wu, Edward B. Cooper, Amy S. Shah, John B. Hogenesch, David F. Smith

**Affiliations:** Division of Human Genetics, Center for Chronobiology, Department of Pediatrics, Cincinnati Children’s Hospital Medical Center, 240 Albert Sabin Way, Cincinnati, OH, 45229; Division of Pulmonary Medicine, Cincinnati Children’s Hospital Medical Center, Cincinnati, OH, 240 Albert Sabin Way, Cincinnati, OH, 45229; Department of Anesthesia, Cincinnati Children’s Hospital Medical Center, 240 Albert Sabin Way, Cincinnati, OH, 45229; Department of Pediatrics, University of Cincinnati College of Medicine, Cincinnati, OH, 240 Albert Sabin Way, Cincinnati, OH, 45229; Division of Endocrinology, Cincinnati Children’s Hospital Medical Center, 3333 Burnet Ave. MLC 7012, Cincinnati, OH, 45229; Divisions of Pediatric Otolaryngology and Pulmonary and Sleep Medicine, Cincinnati Children’s Hospital Medical Center, 3333 Burnet Ave, Cincinnati, OH 45229; Department of Otolaryngology-Head and Neck Surgery, University of Cincinnati School of Medicine, 231 Albert Sabin Way, Cincinnati, OH, 45267

## Abstract

Hospitals operate 24 hours a day, and it is assumed that critical decisions occur continuously around the clock. However, many aspects of hospital operation occur at particular times of day, including medical team rounding and shift changes. It is unclear if this impacts patient care, as an empirical account of 24 h treatment patterns is lacking. We analyzed the daily distribution of ~120K doses of 12 separate drugs in 1,486 inpatients at a major children’s hospital in the U.S. Treatment orders and administration were strongly time-of-day-dependent, marked by distinct morning time surges and overnight lulls. These 24 h rhythms in treatment were remarkably consistent across drugs, diagnoses, and hospital units. In sum, nearly one-third of all 116,975 orders for treatment were placed between 8 AM and 12 PM. This rhythm in hospital medicine coincided with medical team rounding time, not necessarily immediate medical need. Lastly, we show that the clinical response to hydralazine, an acute antihypertensive, is dosing time-dependent and greatest at night, when the fewest doses were administered. The prevailing dogma is that hospital treatment is administered as needed regardless of time of day. Our findings challenge this notion and reveal a potential operational barrier to best clinical care.

**SIGNIFICANCE STATEMENT:** The order and administration of hospital treatment was characterized by morning time surges and overnight lulls, regardless of drug type, diagnosis or care unit. As the first large-scale account of 24 h rhythms in hospital medicine, this study identifies a potential operational barrier to best clinical care. Critical clinical decisions should be made around the clock; pain, infection, hypertensive crisis, and other conditions do not occur selectively in the morning. Systemic bias in the timing of medicine is also at odds with circadian biology, which can influence when certain treatments are most effective or safe. Prevailing dogma is that hospital treatment is administered as needed regardless of time of day. Our findings challenge this notion and suggest that time of day in hospital operations deserves further consideration.

## INTRODUCTION

Hospitals operate 24 hours a day year long. Caretakers are available around the clock, and it is assumed that critical treatment decisions occur continuously regardless of time of day.

However, many universal aspects of hospital operation occur at particular times of day, including medical team rounding and staff shift changes (1, 2). It is unclear if this impacts patient care, as an empirical account of 24 h treatment patterns is lacking.

We conducted an observational cohort study to investigate whether the ordering or administration of hospital treatment is influenced by time of day. Blood pressure (BP) is a continuous phenotype universally tracked in an intensive care unit (ICU) setting. We analyzed the 24 h distribution of order and administration times in 1,481 patients who received at least one dose of hydralazine, a commonly used antihypertensive in the ICU, at a tertiary pediatric hospital from 2010 to 2017. Although hydralazine treatment was the basis for patient inclusion, our analysis also included 11 other drugs commonly administered to these patients.

We found that treatment was strongly time-of-day-dependent, characterized by distinct morning time surges and overnight lulls. Leveraging this large dataset of ~ 120K doses, we examined the 24 h distribution of order and administration times of multiple different drug classes, across different care units. Lastly, we explored the impact of dosing time on clinical outcomes.

## RESULTS

Hydralazine is a vasodilator used to treat essential hypertension. We analyzed its daily use in 1,481 patients who received at least one intravenous (IV) dose of the drug. In total, 5,294 orders for IV hydralazine were placed, each stipulating a dosing frequency of “once”, “as needed”, or “scheduled” (Table 1). We tracked both order time (when a care provider replaces an electronic request for a drug) and dosing time (when the patient receives the drug) for each patient. Initial analyses considered only the first dose administered after each order (Methods).

**Table 1.** Patient and treatment summary.

| Patient admissions (n = 7,694) |  |  |  |  |  |  |
| --- | --- | --- | --- | --- | --- | --- |
| Age (y) | 1–5 | 5–7 | 7–10 | 10–15 | 15–20 | > 20 |
| **Sex (%)** |  |  |  |  |  |  |
| *Female* | 241 (40) | 251 (42) | 596 (39) | 695 (41) | 507 (42) | 1068 (52) |
| *Male* | 359 (60) | 347 (58) | 934 (61) | 1018 (59) | 691 (58) | 987 (48) |
| **Diagnosis %** |  |  |  |  |  |  |
| *Fever* | 8 | 11 | 12 | 12 | 6 | 5 |
| *Respiratory distress* | 3 | 3 | 3 | 3 | 2 | 3 |
| *Hemophagocytic syndromes* | 2 | 2 | 2 | 1 | 1 | 1 |
| *Leukemia* | 2 | 3 | 2 | 5 | 4 | 4 |
| *Dehydration* | 1 |  | 2 | 1 | 1 | 1 |
| *Neutropenia* | 1 | 2 | 2 | 2 | 2 | 1 |
| *Renal failure / injury* | 1 | 1 | 3 | 3 | 1 | 3 |
| *Pneumonia* | 1 | 1 | 2 | 2 | 2 | 3 |
| *Cystic fibrosis* | < 1 | < 1 | < 1 | < 1 | 1 | 3 |
| *Other* | 30 | 32 | 39 | 37 | 42 | 49 |
| *Not reported* | 51 | 44 | 33 | 34 | 38 | 27 |

| Treatment orders (n = 116, 975) and doses (n = 590,196) |  |  |  |  |  |  |  |  |  |
| --- | --- | --- | --- | --- | --- | --- | --- | --- | --- |
| Drug | Orders | % Order Frequency |  |  | Doses |  | % Treatment Unit |  |  |
|  |  | Once | Sched. | As needed | Total | % First | ICU | Floor | Other /NA |
| *Hydralazine* | 5,599 | 42 | 6 | 52 | 17,760 | 32 | 43 | 38 | 19 |
| *Acetaminophen* | 22,885 | 30 | 14 | 55 | 88,794 | 26 | 22 | 54 | 25 |
| *Furosemide* | 18,370 | 65 | 35 | 0 | 53,194 | 35 | 43 | 42 | 15 |
| *Morphine* | 14,885 | 34 | 58 | 9 | 71,370 | 21 | 49 | 31 | 20 |
| *Lorazepam* | 11,442 | 28 | 40 | 31 | 97,177 | 12 | 44 | 35 | 21 |
| *Ondansetron* | 8,701 | 19 | 40 | 41 | 74,537 | 12 | 14 | 61 | 25 |
| *Fentanyl* | 8,564 | 49 | 2 | 49 | 28,655 | 30 | 69 | 7 | 24 |
| *Hydrocortisone* | 8,413 | 22 | 72 | 5 | 60,815 | 14 | 25 | 57 | 18 |
| *Diphenhydramine* | 7,998 | 39 | 12 | 49 | 28,946 | 28 | 12 | 64 | 24 |
| *Methylprednisolone* | 4,905 | 28 | 72 | 1 | 28,353 | 17 | 36 | 43 | 21 |
| *Labetalol* | 2,870 | 29 | 46 | 31 | 22,550 | 13 | 40 | 38 | 22 |
| *Vancomycin* | 2,343 | 16 | 84 | 0 | 18,145 | 13 | 30 | 33 | 37 |

Hydralazine order and first dosing times were non-uniformly distributed over 24 h (Kuiper’s test, *P* < 0.01), marked by distinct morning-time surges and overnight lulls (Fig. 1A). Nearly twice as many treatments were ordered (1,495) and dosed (1,445) between 10 AM to 4 PM compared to 2 AM–8 AM (783 and 899, respectively). The profiles were described by 24 h rhythms by three separate detection methods (cosinor analysis (3), JTK_CYCLE (4), and RAIN (5), *P* < 0.05) (Fig. 1B, *SI Appendix, Table 1*). These rhythms were consistent across patient age groups and treatment units—ICU or general floor (Fig. 2).

**Fig 1.**
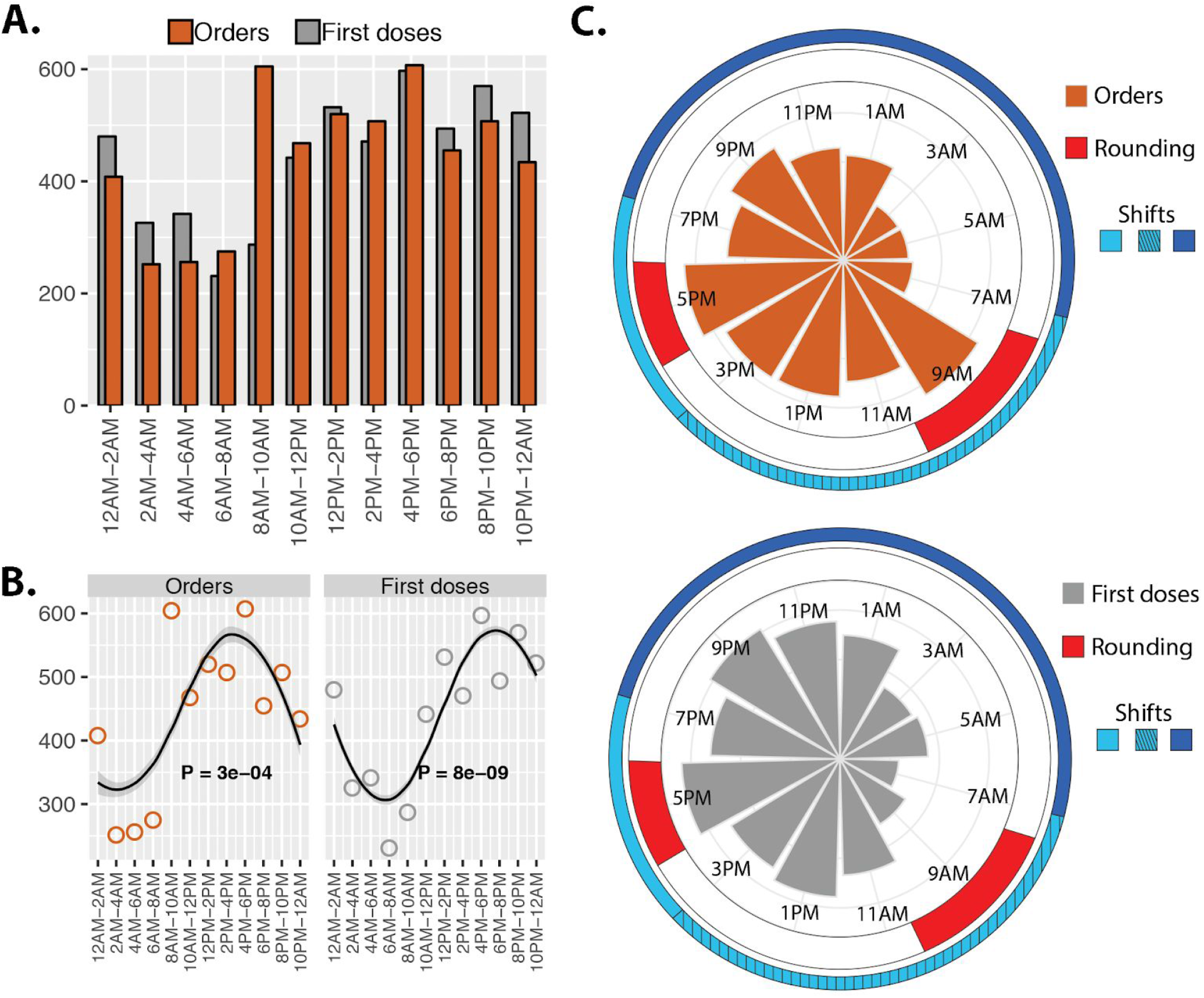
Daily rhythms in the timing of hydralazine treatment. **(A)** Nearly twice as many treatments were ordered (1,495) and administered (1,445) 10 AM–4 PM compared to 2 AM–8 AM (783 and 899 respectively). **(B)** The number of orders and first doses in 2-hour bins were modeled by a cosinor waveform with a 24 h period (cosinor analysis, *P* < .05). **(C)** Polar histograms indicate the times of hydralazine orders (top) and first doses (bottom). Team rounds occur from 7.30 AM–10.30 AM and 4 PM–6 PM. Clinical caretakers at this study site work either a 7 AM–7 PM (light blue), 7 AM–3 PM (light blue hashed), or 7 PM–7 AM shift (dark blue). This schedule is consistent throughout hospitals in the U.S.

To determine if the 24 h rhythm in dosing of hydralazine reflected immediate medical need for treatment, we analyzed the systolic BP recording immediately prior to drug dosing (d-SBP) (Fig 2A). d-SBPs were highly variable within each 2 h time bin. There were no statistically discernible differences in d-SBP between time bins (one-way ANOVA, *P* > 0.1), suggesting that the 24 h rhythms in treatment were not solely driven by immediate medical need. Why is the distribution of BPs discordant with dosing time?

**Fig 2.**
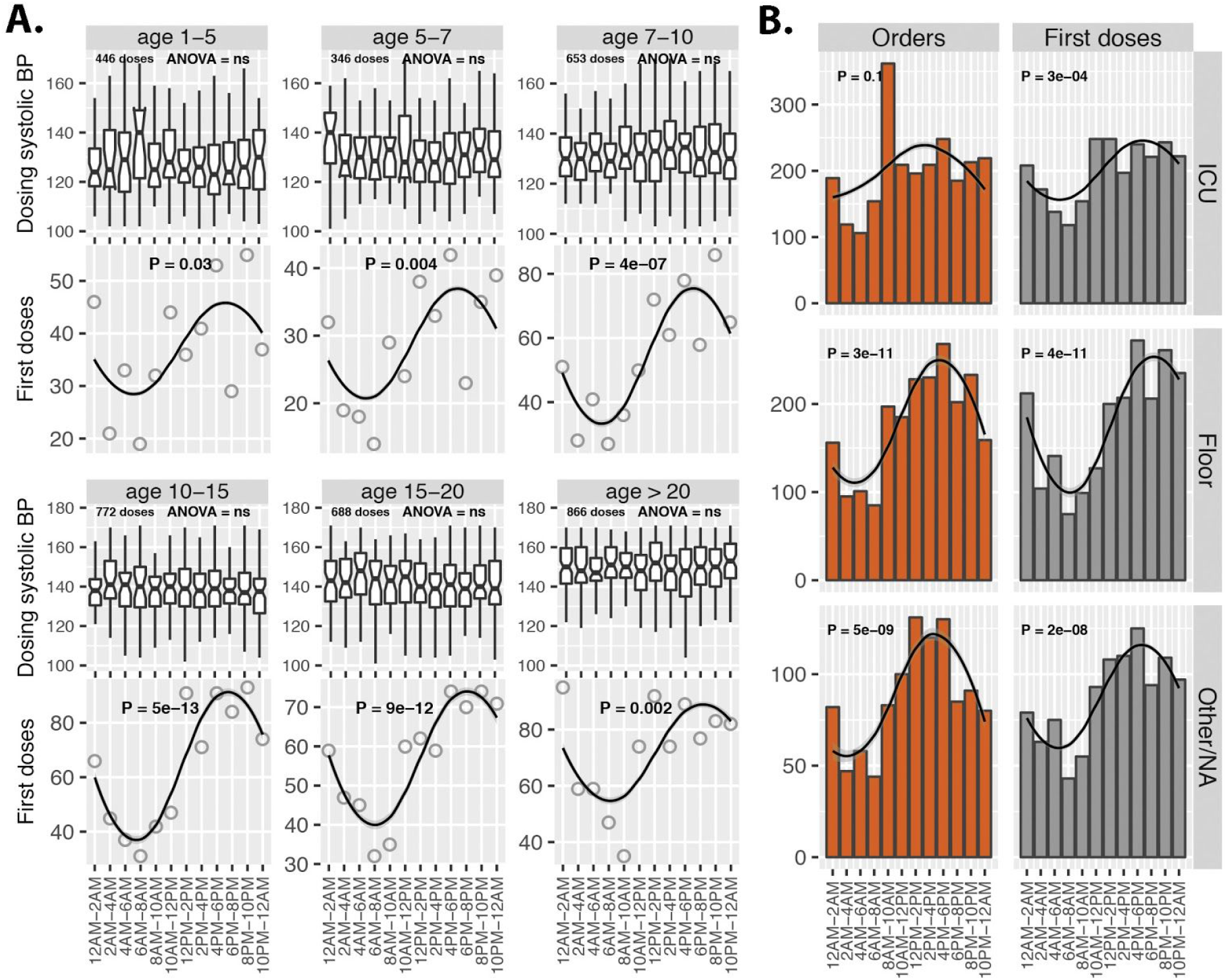
Rhythms in hydralazine orders and first doses across patient ages and hospital units. **(A)** Top panels: Systolic BP immediately prior to hydralazine dosing (d-SBP). d-SBPs were assigned to time bins according to time of hydralazine administration. Within each age group, no differences in d-SBP were detected between time bins (one-way ANOVA, P > .05). Boxplot center lines indicate median d-SBP, whiskers the upper and lower 25th percentiles. Outliers, defined as 1.5x the interquartile range, are not shown. Bottom panels: The number of hydralazine first doses in each time bin modeled by a cosinor waveform with a 24 h period. **(B)** 24 h rhythms in hydralazine treatment (cosinor regression) across treatment units.

The morning surge in hydralazine order times coincide with team rounding and a medical staff shift change (Fig. 1C). Caretaker shifts at Cincinnati Children’s Hospital Medical Center (CCHMC) include a 7 AM–7 PM, 7 AM–3 PM, or 7 PM–7 AM shifts. Team rounds typically occur from 7.30 AM-10.30 AM and 4 PM-6 PM. These schedules are consistent across the U.S.

We next tested if the 24 h patterns in hydralazine use generalized to other therapies, including analgesics, anti-infectives, antihistamines, diuretics, and other drug classes commonly used in the hospital. For each of the additional 11 drugs analyzed, order and first-dose times were non-uniformly distributed (Kuiper’s test, P < 0.01) with morning surges and overnight lulls (Fig. 3A, *SI Appendix*, *Fig. S*1A). The majority (9/11) were described by 24 h rhythms (cosinor, *P* < 0.05; RAIN or JTK_CYCLE, *P* < 0.05), regardless of treatment unit (*SI Appendix, Table 1 and* *Fig. S*1B). Overall, nearly one-third of the total 116,975 drug orders were placed during the 4 h time window from 8 AM to 12 PM. This morning surge in ordering of all drugs coincides with team rounding (Fig. 3B & 3C).

Our previous analyses included all order frequencies (once, scheduled, or as needed) but considered only the first dosing time from each order. Next, we analyzed scheduled orders only (e.g., “take every 8 h”), focusing on hydralazine as an example (*SI Appendix*, *Fig. S*2). As expected, the first dosing times were rhythmically distributed. All subsequent dosing times, however, were uniformly distributed over 24 h, without discernible rhythms (cosinor analysis, *P* > 0.2). Time-of-day bias in the original ordering time, therefore, impacts all subsequent dosing times.

**Fig 3.**
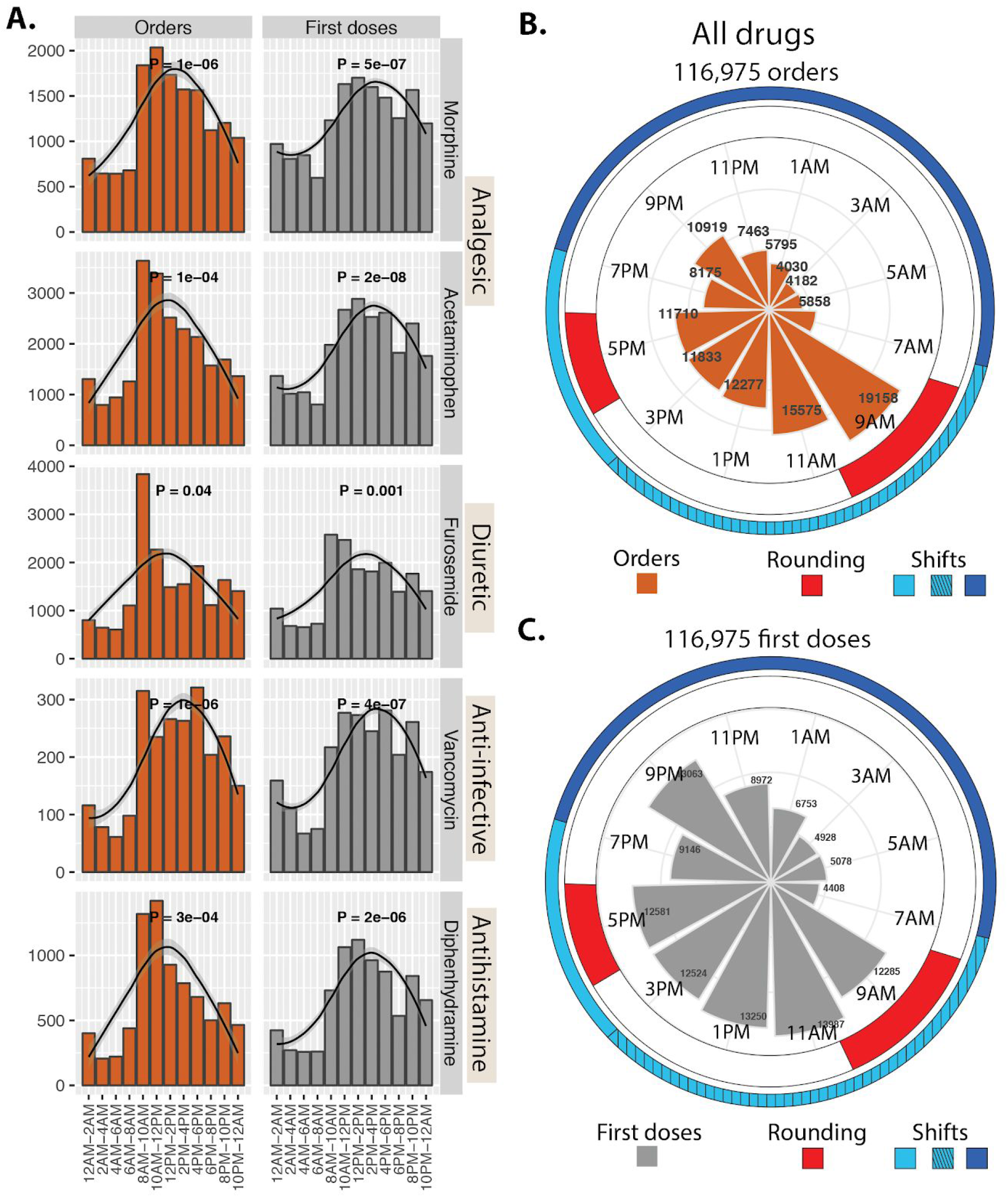
Rhythms in treatment across drug classes coincide with hospital-wide operational activity. **(A)** Ordering and administration of several different drug classes all showed 24 h rhythms marked by morning time surges and nighttime lulls. **(B)** The polar histogram indicates the times of all drug orders and **(C)** first doses administered. The onset of peak ordering time coincides with morning team rounding (indicated in red). Clinical caretakers at this study site (CCHMC) work either a 7 AM–7 PM (light blue), 7 AM–3 PM (light blue hashed), or 7 PM–7 AM shift (dark blue). Team rounds occur from 7.30 AM–10.30 AM and 4 PM–6 PM. This schedule is consistent throughout hospitals in the US.

Rhythms in hospital medicine may conflict with circadian biology, as dosing time influences responses to many types of treatment (6). For example, in the outpatient setting, short-acting antihypertensives are most effective at lowering BP if taken before bedtime (7). To test if this applies to acute antihypertensive therapy in the hospital, we analyzed inpatient responses to 7,132 doses of hydralazine as a function of dosing time (Fig. 4, Methods). Response was computed as the percent change between dosing BP (just before dose) and mean BP over the 3 h following each dose. To control for the impact of dosage, we stratified doses by concentration (mg/kg body weight) (Fig. 4A). The response to hydralazine varied over 24 h (ANCOVA, diastolic P = 3e-11; systolic P = 0.002) in the most common dosage group (0.1 ≥ mg/kg ≤ 0.2; n = 5,188 of 7,132 doses) (Figs. 4B and C). BP was most responsive to nighttime dosing (10 PM–2 AM), and least responsive to morning (6 AM–10 AM) and late afternoon (2 PM − 6 PM) dosing (Tukey’s test, P-adjusted < 0.001; see *SI Appendix, Table 2* for difference and confidence levels for pairwise comparisons). Both diastolic and systolic responses followed this pattern, although the variation in diastolic BP was more pronounced. In sum, nighttime hydralazine was associated with an ~ 3–4% greater reduction in diastolic BP on average than morning dosing (Tukey’s test, difference = −3.4, 95% CI upper level = −5.5, lower level = −1.4, P-adjusted = 2e-06). We did not detect 24 h variation in response for any of the other three dosage groups, which included far fewer doses (*SI Appendix*, *Fig. S*3). Given the many potential sources of variation in response in this cohort, a sufficiently large sample may be required to detect 24 patterns. Nevertheless, for the majority of doses, the clinical response to hydralazine varied over 24 h.

**Fig 4.**
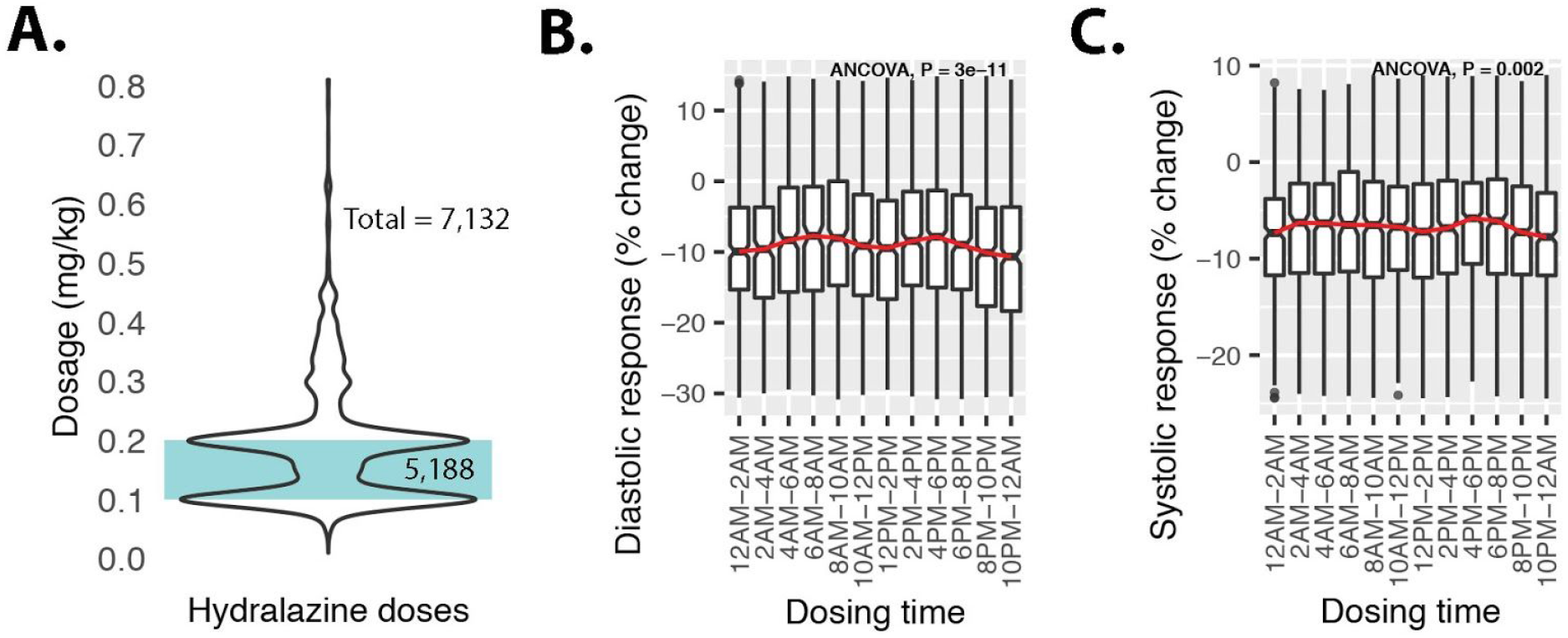
Clinical response to hydralazine varies by time of administration. **(A)** 5,188 doses of hydralazine ranging from 0.1 to 0.2 mg/kg body weight. Percent change in **(B)** diastolic and **(C)** systolic BP as a function of dosing time. Percent change was computed as the difference between dosing BP (just before dose) and mean BP over the 3 h following each dose. Response to hydralazine varied over 24 h (ANCOVA, diastolic P = 3e-11 and systolic P = 0.002) (see Methods). Boxplot center lines indicate median, whiskers the upper and lower 25th percentiles. Outliers were defined as more than 1.5x the interquartile range (black circles). For visual aid, a line was fit through the medians. Post-hoc analyses (Tukey’s test) identified larger responses in early afternoon (12 PM–2 PM) and nighttime (10 PM – 12 AM) compared to the morning (8 AM – 10AM) and late afternoon (4PM – 6PM) (see *SI Appendix, Table* 2).

## DISCUSSION

Order and first dosing times of antihypertensives, analgesics, anti-infectives, and other drug therapies all followed a 24 h rhythm characterized by a morning surge and an overnight lull. There are no practice guidelines or institutional policies specifying time-of-day recommendations for any of the drugs in this study. The surge in orders (8 AM–10 AM) coincides with rounding – when the medical care team visits each patient and collaboratively develops a treatment plan. The majority of orders for diagnostics, therapies, and referrals to specialty services are typically placed during rounds or immediately thereafter, once final consensus is reached.

Systemic bias towards treatment at a particular time of day is problematic. Anti-infectives should be administered when an infection is identified, whereas analgesics should be administered when there is pain. Neither of these happen selectively in the morning. In fact, evidence shows that pain is more severe in the evening (8). Circadian clocks coordinate physiologic functions (9) and affect therapeutic responses according to the time of day (6). Short-acting antihypertensives taken by millions of adults, for example, are most effective before bedtime (10). Here, we show that this is also true in the pediatric hospital setting. The clinical response to an acute antihypertensive, hydralazine, varied by 3–4% over 24 h. Paradoxically, patients were more responsive to therapy during the window of time when the fewest doses were ordered and administered.

Conversely, other treatments have shown to be more effective if administered in the morning (vaccines (11), antipsychotics (12)), or midday (corticosteroids (13), radiotherapy (14), cardiac surgery (15)). There is no single optimal dosing time for all drugs and patients.

Hospital medical staff are available to provide treatment around the clock, and prevailing dogma is that treatment is given as needed regardless of time of day. Our findings challenge this notion and reveal a potential barrier to best clinical care. Interestingly, studies show that more frequent rounding by nurses can improve measures of patient care (16). Would medical rounding designed to be more evenly distributed over 24 h improve clinical outcomes? Indeed, there are other long-standing aspects of hospital operation that have been challenged. For example, recent evidence shows that traditional continuous dim lighting in ICUs (17) lacks a medical rationale, disrupts patients, and may impede recovery (18–21).

As with all retrospective clinical studies, there were a number of potential sources of variability whose specific contribution could not be evaluated. For example, patient genotype (SNPs can affect hydralazine metabolism (22)), drug–drug interactions, or special clinical circumstances could each impact the clinical response to hydralazine. We hope that future multicenter prospective trials will further elucidate the importance of drug timing on patient outcomes. In sum, time of day in hospital operations deserves further consideration.

## METHODS

### Study design

We performed an observational cohort study of 1,481 patients receiving hydralazine, a commonly used antihypertensive in the ICU, at a tertiary pediatric hospital from Jan 2010 to Dec 2017. All patients receiving hydralazine while admitted to the hospital were included. Demographic data, medical history, BP, and order and administration times of 12 different drug therapies were recorded. Institutional review board (IRB) approval was obtained prior to initiation of this project.

We analyzed the daily distribution of order and administration times in 1,481 patients who received at least one dose of hydralazine. Although hydralazine treatment was the basis for patient inclusion, our analysis also included 11 other drugs commonly administered to these patients. Patients were diverse in age, sex, race, and diagnoses (Table 1). Treatments were administered in ICUs or general floors. Orders for treatment were categorized as “once”, “as needed”, or “scheduled”.

### Statistical analyses — tests for uniformity and periodic pattern of orders and doses

For each drug, we tested if 24 h patterns in ordering or administration deviated from a uniform distribution (Kuiper’s test (23)). To avoid bias from scheduled doses (“take every 8 h”, for example), we limited analyses to the first dose after each order unless otherwise specified. To test for 24 h rhythms in treatment, we applied three independent detection methods, cosinor analysis (3), JTK_CYCLE (4), and RAIN (5). For each drug, orders and first-doses were discretized into 2-hour time bins. The amplitude, phase, and overall significance of each rhythmic model is presented in *SI Appendix, Table 1*.

### Test for daily variation in dosing BP

To analyze BPs associated with hydralazine administration, we identified the BP recording immediately prior to each dose, referred to as the dosing systolic BP (d-SBP). Doses were excluded from analysis if the d-SBP was more than 1 h prior to drug administration.

### Test for daily variation in response to hydralazine

We analyzed inpatient responses to hydralazine as a function of dosing time. Response was defined as the percent change between dosing BP (just before dose) and mean BP over the 3 h following each dose. BP was measured irregularly over time, so we required at least 3 BP records in the 3 h response window for a dose to be considered. We eliminated outlier responses, defined as greater than the 97th or less than the 3rd percentiles of all dose responses. To control for the impact of dosage on response, we stratified doses by concentration (mg/kg body weight) into 4 dosage subgroups, each with at least 300 doses. To test for variation in response over 24 h, we applied a linear model (ANOVA) with dosing BP as a covariate to control for effect of different starting BP. We ran a post-hoc Tukey’s test to estimate the mean difference in response, 95th percentile confidence levels, and multiple hypothesis adjusted P-values for all possible pairwise comparisons between time bins.

## SUPPLEMENTAL INFORMATION

**Fig S1.**
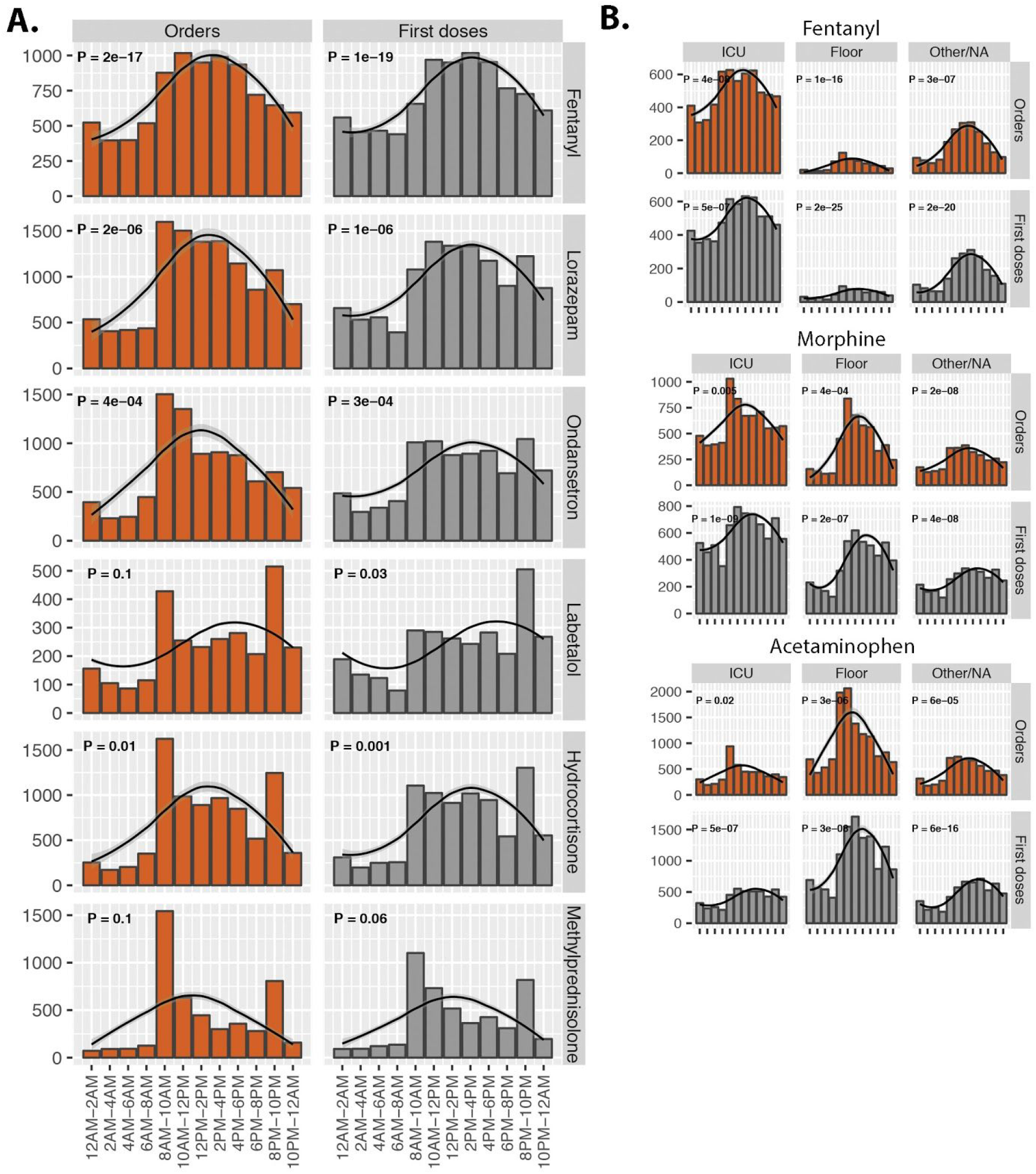
Rhythms in orders and first doses across drug classes and hospital units. (A) Orders and first doses of different drug classes were all non-uniformly distributed over 24 h, the majority with detectable 24 h rhythms (cosinor analysis, P < 0.05, see also *SI Appendix, Table 1*). (B) Daily rhythms in orders and first doses were independent of the hospital treatment unit.

**Fig S2.**
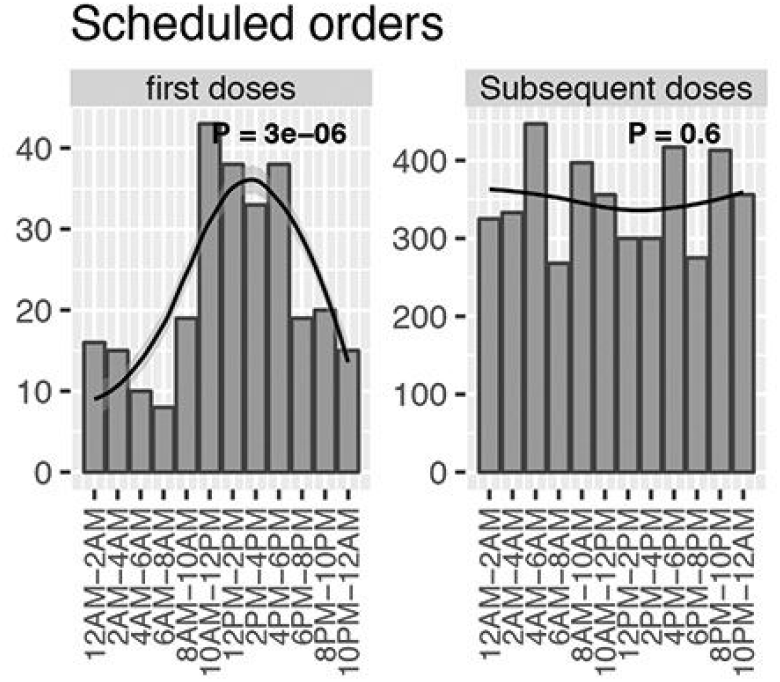
Scheduled-only orders for hydralazine. Scheduled orders (eg., take every 8 hrs) for hydralazine. First doses were described by a 24 h rhythm, whereas all subsequent doses were more uniformly distributed over 24 h and without a statistically discernible rhythm (P = 0.6).

**Fig S3.**
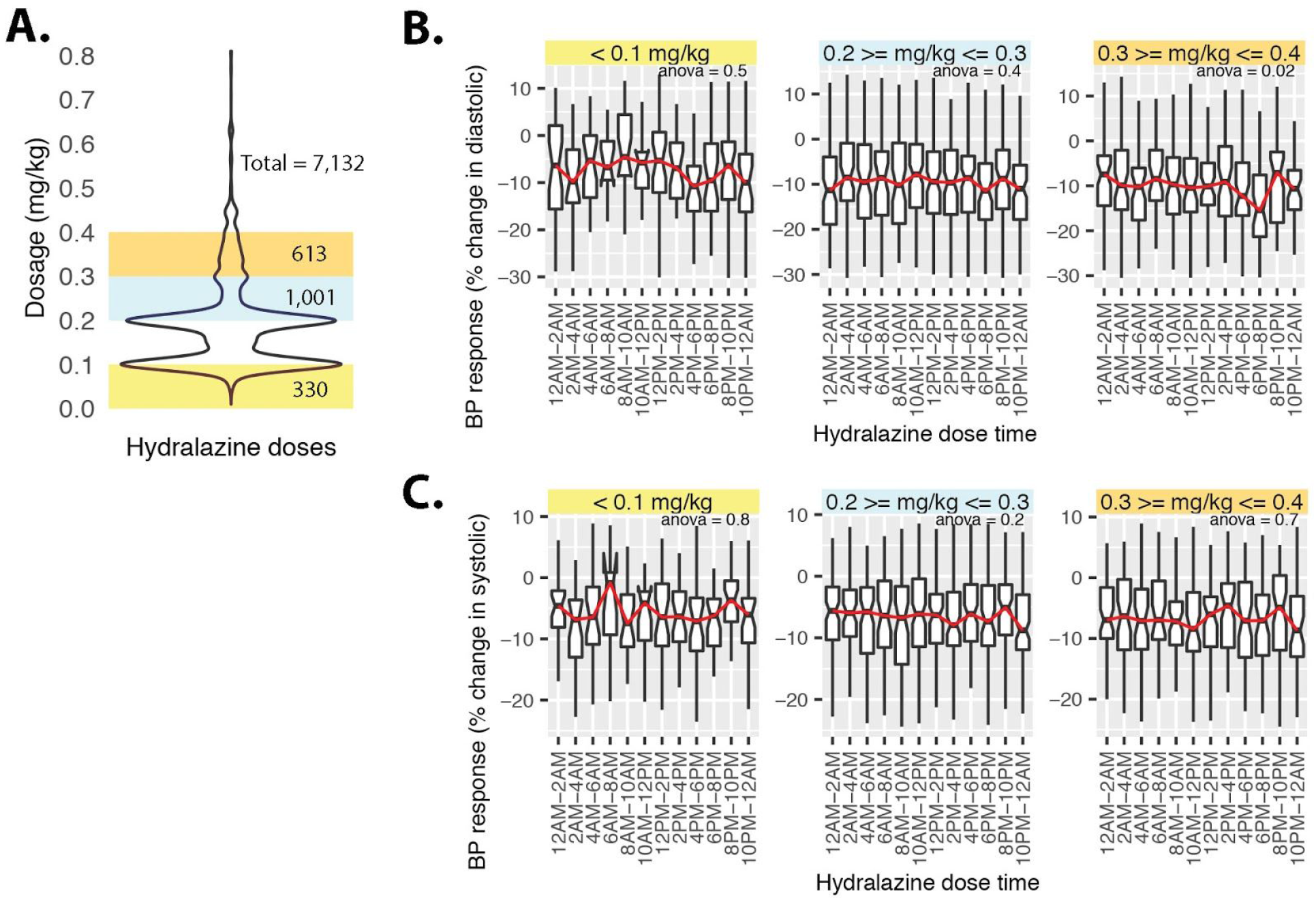
Clinical response to hydralazine by time of administration. **(A)** Hydralazine at dosages ranging from 0–0.1, 0.2–0.3, and 0.3–0.4mg/kg body weight. Percent change in **(B)** diastolic and **(C)** systolic BP as a function of dosing time. Percent change was computed as the difference between dosing BP (just before dose) and mean BP over the 3 h following each dose. Boxplot center lines indicate median response, whiskers the upper and lower 25th percentiles. Outliers, defined as more than 1.5x the interquartile range, are shown as black circles. For visual aid, a line was fit through the medians.

**Table S1.** Detection of 24 h rhythms in orders and first doses. Results from two independent tests, RAIN and JTK_CYCLE, for periodic patterns in data. For each drug, orders and first-doses were discretized into 2-hour time bins. Peak phase and overall significance were computed for both models specifying a period of 24 h.

|  | RAIN |  |  | JTK |  |  |  |
| --- | --- | --- | --- | --- | --- | --- | --- |
|  | pVal | peak phase | period | pVal | period | peak phase | amplitude |
| Acetaminophen_doses | 3.50E-04 | 18 | 24 | 0.02 | 24 | 17 | 884.4 |
| Acetaminophen_orders | 1.20E-05 | 18 | 24 | 0.06 | 24 | 16 | 872.6 |
| Diphenhydramine_doses | 1.40E-04 | 16 | 24 | 0.02 | 24 | 17 | 338.4 |
| Diphenhydramine_orders | 2.40E-05 | 16 | 24 | 0.02 | 24 | 16 | 341.2 |
| Fentanyl_doses | 1.50E-04 | 16 | 24 | 0 | 24 | 17 | 254.9 |
| Fentanyl_orders | 2.20E-05 | 16 | 24 | 0 | 24 | 16 | 293.1 |
| Furosemide_doses | 2.50E-03 | 18 | 24 | 0.25 | 24 | 18 | 533.2 |
| Furosemide_orders | 3.80E-02 | 18 | 24 | 0.25 | 24 | 17 | 423.2 |
| Hydralazine_doses | 4.10E-03 | 14 | 24 | 0.01 | 24 | 20 | 143 |
| Hydralazine_orders | 4.20E-03 | 10 | 24 | 0.01 | 24 | 18 | 126.6 |
| Hydrocortisone_doses | 1.10E-01 | 18 | 24 | 0.25 | 24 | 17 | 430.6 |
| Hydrocortisone_orders | 1.30E-02 | 18 | 24 | 0.17 | 24 | 16 | 439.1 |
| Labetalol_doses | 4.70E-01 | 18 | 24 | 1 | 24 | 18 | 67.2 |
| Labetalol_orders | 1.00E-01 | 12 | 24 | 0.11 | 24 | 18 | 72.8 |
| Lorazepam_doses | 7.50E-04 | 18 | 24 | 0.04 | 24 | 17 | 434.9 |
| Lorazepam_orders | 8.20E-05 | 18 | 24 | 0.04 | 24 | 16 | 570.5 |
| Methylprednisolone_doses | 1.60E-02 | 16 | 24 | 0.17 | 24 | 15 | 236.2 |
| Methylprednisolone_orders | 1.60E-02 | 16 | 24 | 0.17 | 24 | 15 | 215 |
| Morphine_doses | 4.10E-04 | 18 | 24 | 0.01 | 24 | 17 | 453.6 |
| Morphine_orders | 1.20E-05 | 18 | 24 | 0.06 | 24 | 16 | 604.2 |
| Ondansetron_doses | 9.10E-02 | 18 | 24 | 0.25 | 24 | 18 | 286.9 |
| Ondansetron_orders | 8.10E-05 | 18 | 24 | 0.02 | 24 | 16 | 394.9 |
| Vancomycin_doses | 2.50E-03 | 18 | 24 | 0.02 | 24 | 18 | 90.5 |
| Vancomycin_orders | 6.10E-03 | 16 | 24 | 0.01 | 24 | 18 | 98.1 |

## Author Contributions

Drs Ruben and Smith had full access to all of the data in the study and take responsibility for the integrity of the data and the accuracy of the data analysis. *Concept and design:* Ruben, Francey, Smith, Hogenesch. *Acquisition, analysis, or interpretation of data:*All authors. *Drafting of the manuscript:* Ruben, Smith. *Critical revision of the manuscript for important intellectual content:* All authors. *Statistical analysis:* Ruben, Wu, Hogenesch. *Administrative, technical, or material support:* Ruben, Wu, Guo, Hogenesch. *Supervision:* Ruben, Smith, Hogenesch

